# Implementation of residue-level coarse-grained models in GENESIS for large-scale molecular dynamics simulations

**DOI:** 10.1101/2021.10.21.465249

**Authors:** Cheng Tan, Jaewoon Jung, Chigusa Kobayashi, Diego Ugarte La Torre, Shoji Takada, Yuji Sugita

**Author notes:** Corresponding author, (YS).

## Abstract

Residue-level coarse-grained (CG) models have become one of the most popular tools in biomolecular simulations in the trade-off between modeling accuracy and computational efficiency. To investigate large-scale biological phenomena in molecular dynamics (MD) simulations with CG models, unified treatments of proteins and nucleic acids, as well as efficient parallel computations, are indispensable. In the GENESIS MD software, we implement several residue-level CG models, covering structure-based and context-based potentials for both well-folded biomolecules and intrinsically disordered regions. An amino acid residue in protein is represented as a single CG particle centered at the Cα atom position, while a nucleotide in RNA or DNA is modeled with three beads. Then, a single CG particle represents around ten heavy atoms in both proteins and nucleic acids. The input data in CG MD simulations are treated as GROMACS-style input files generated from a newly developed toolbox, GENESIS-CG-tool. To optimize the performance in CG MD simulations, we utilize multiple neighbor lists, each of which is attached to a different nonbonded interaction potential in the cell-linked list method. We found that random number generations for Gaussian distributions in the Langevin thermostat are one of the bottlenecks in CG MD simulations. Therefore, we parallelize the computations with message-passing-interface (MPI) to improve the performance on PC clusters or supercomputers. We simulate Herpes simplex virus (HSV) type 2 B-capsid and chromatin models containing more than 1,000 nucleosomes in GENESIS as examples of large-scale biomolecular simulations with residue-level CG models. This framework extends accessible spatial and temporal scales by multi-scale simulations to study biologically relevant phenomena, such as genome-scale chromatin folding or phase-separated membrane-less condensations.

**Author summary:** Molecular dynamics (MD) simulations have been widely used to investigate biological phenomena that are difficult to study only with experiments. Since all-atom MD simulations of large biomolecular complexes are computationally expensive, coarse-grained (CG) models based on different approximations and interaction potentials have been developed so far. There are two practical issues in biological MD simulations with CG models. The first issue is the input file generations of highly heterogeneous systems. In contrast to well-established all-atom models, specific features are introduced in each CG model, making it difficult to generate input data for the systems containing different types of biomolecules. The second issue is how to improve the computational performance in CG MD simulations of heterogeneous biological systems. Here, we introduce a user-friendly toolbox to generate input files of residue-level CG models containing folded and disordered proteins, RNAs, and DNAs using a unified format and optimize the performance of CG MD simulations via efficient parallelization in GENESIS software. Our implementation will serve as a framework to develop novel CG models and investigate various biological phenomena in the cell.

## Introduction

Molecular dynamics (MD) simulation of a biomolecule in solution or membrane has been successfully applied to biophysical or biochemical problems at high spatial and temporal resolutions that are unattainable by experimental techniques. Recently, biological phenomena involving many biomolecules at larger scales have attracted much more attention than before in molecular and cellular biology, as well as medical and pharmaceutical studies. Genome-scale chromatin folding [1] and phase-separated membrane-less condensations [2] are well-known examples, which have already been targets of MD simulation studies [3–7]. To obtain biological insights on these phenomena, further methodological and computational challenges need to be met. An enormous number of atoms, for instance, more than 10-100 million atoms, are involved in such target systems, requiring us to improve the computational performance of the simulations. Alongside the size problem, there are also barriers to achieving sufficient conformational sampling in biologically meaningful time scales. Considering these issues, coarse-grained (CG) models of biomolecules are very useful in MD simulations, being a trade-off between modeling accuracy and computational efficiency. The CG models are able to decrease the number of degrees of freedom in a system and to smooth out the ruggedness of free energy landscapes, both of which efficiently accelerate MD simulations of biological systems [8].

Although there are various levels of coarse-graining [9–13], they can be categorized roughly into two classes of models in terms of their theories and parameterizations: physical-based or structure-based CG models. The MARTINI model [13] is one of the representatives in the former, while the Gō model [14–16], which describes the funnel-like energy landscape of protein folding, is probably the most well-known structure-based one. The resolution of each CG model, namely, how many heavy atoms are integrated as a single CG particle, is another crucial point in the choice of CG models. We focus on residue-level CG models [17], whose interaction potentials are primarily described via structure-based ones. These models have been utilized in many simulation studies of folding dynamics of single proteins [14], conformational dynamics of nucleic acids [18,19], and complex molecular machinery [20]. In these models, an amino acid residue is usually simplified as one CG particle on the *C_α_* atom, and a nucleotide is represented using three particles corresponding to the phosphate (P), sugar (S), and base (B), respectively (Fig 1). With this representation, proteins and nucleic acids are modeled with a similar resolution of around ten heavy atoms per single CG particle. Furthermore, as an extension from the original Gō model, the atomic interaction-based CG model version 2+ (AICG2+) [21] protein model introduces flexible local potentials and sequence-dependent contact parameters to gain a finer description of the free energy landscapes. The same strategy has been applied to RNA molecules [22]. On the other hand, for DNA, the 3SPN.2C model has been designed to describe the sequence-dependent geometric and mechanical properties of double-stranded DNA (dsDNA) [23].

**Fig 1.**
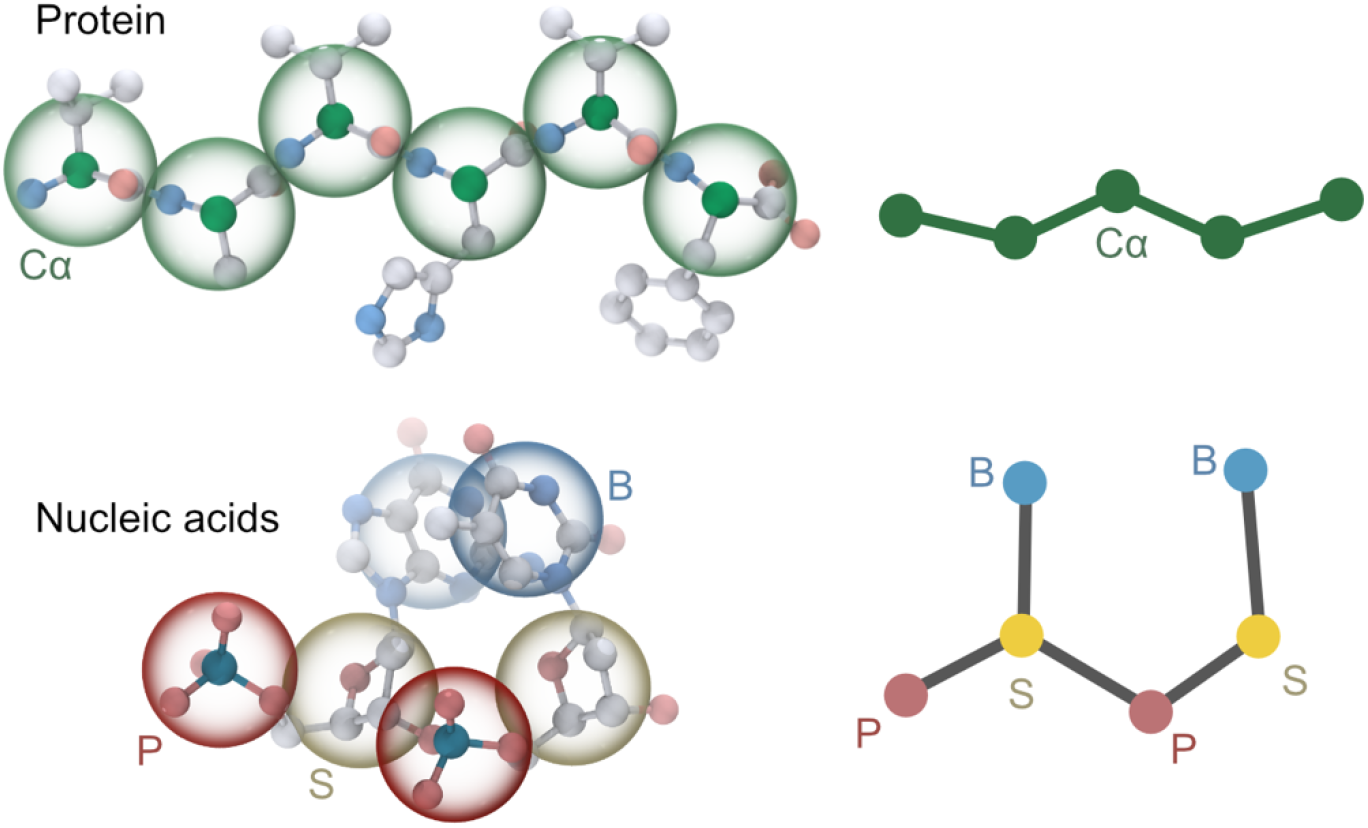
Mapping of a protein (top) and nucleic acids (bottom) from atomistic structures to the residue-level CG models. Each amino acid residue is represented with one CG particle (C_α_), and each nucleotide is represented using three CG particles, phosphate (P), sugar (S), and base (B), respectively. Left: three-dimensional structure of protein and nucleic acids. Big and transparent spheres are the CG particles, with C_α_ in green, P in red, S in yellow, and B in blue. Small and solid spheres represent heavy atoms, with phosphorus in dark blue, nitrogen in pale blue, oxygen in red, C_α_ in green, and all the other carbon in white. Right: two-dimensional ball-and-stick cartoons of protein and nucleic acids abstracted from the left structures, using the same color scheme.

The AICG2+ and 3SPN.2C models have been used jointly in simulations of protein-DNA complexes [24–26], although these two were developed by different research groups. In these simulations, electrostatic and excluded volume interactions between proteins or between proteins and nucleic acids are necessary for heterogeneous biological systems. In addition to generic and physical interactions, the PWMcos method integrates high-throughput sequencing experimental results in the form of the position weight matrix (PWM) with complex structural information to describe the recognition of DNA bases by proteins [27]. Not only for the well-defined folded structures of proteins, residue-level CG models can also be used to study highly flexible and intrinsically disordered regions (IDR). In particular, two pairwise interaction models, Hydrophobicity scale (HPS) [28] and Kim-Hummer (KH) [29], have been tested and refined to reproduce the phase behaviors of protein IDRs [30,31] and RNAs [32] by Best, Mittal, and their coworkers.

The CG models mentioned above were implemented in general-purpose MD packages such as LAMMPS [33,34], GROMACS [35,36], NAMD [35,37], and OpenMM [38], or specialized CG software such as CafeMol [39]. Although these implementations are convenient tools for performing CG simulations, there are still some barriers for regular users or even developers. Each residue-based CG model is usually developed by different groups in different MD packages. If someone wants to use two of them simultaneously, it is not straightforward to combine two different CG models by treating their input or output data properly. Therefore, there is a necessity to incorporate the latest CG models into a unified working environment and provide a user-friendly tool for preparing MD input files. Furthermore, simultaneous usages of different CG models can be a powerful tool in the cutting-edge studies of protein-nucleic acid complexes in cellular environments. For example, the liquid-liquid phase-separated (LLPS) systems often involve complicated interactions among multi-domain proteins [40], IDRs [41], RNAs [42], and DNAs [43] and reach the length scales of hundreds to thousands of nanometers [44]. Such systems require both proper modeling of biomolecular interactions and efficient handling of computational resources.

In the current work, we present an implementation of several residue-level CG models of protein and nucleic acids in the GENESIS MD software [45,46]. In conjunction, we provide a collection of Julia [47] scripts, which we call GENESIS-CG-tool, to help users generate CG MD input files of complex biomolecular systems. The optimization and parallelization of CG MD simulations are the other focus in the current work. Although all-atom MD simulations in GENESIS have been highly optimized and parallelized, additional efforts are required to improve the performance of CG MD simulations because of the uniqueness of individual interaction potentials. We also noticed that some computations that take negligible fractions of time in all-atom MD simulations could be a bottleneck in CG MD simulations. After the implementation in GENESIS, memory and performance benchmark tests are carried out. CG MD simulations in GENESIS are applied to several cases of different sizes and molecular compositions, such as protein diffusion on DNA and LLPS consisting of protein IDRs and RNAs. The atomistic and CG MD simulations available in GENESIS can be a practical framework to investigate cellular-scale biological phenomena at multi-scale resolutions.

## Design and implementation

### Basic interaction terms

We first define basic interaction terms that are commonly used in various residue-level CG models. Each model, such as AICG2+, 3SPN.2P, HPS/KH, combines several basic interaction terms to represent intra- and inter-molecular potentials of proteins and/or nucleic acids.

### Bond potential

The most typical bond potential has the following harmonic form:

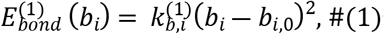

where *b_i_* is the bond length between two neighboring CG particles, *b*_*i*,0_ is the value of *b_i_* in the reference structure, and 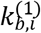 is the force constant.

A higher-order term is also used in some CG models, such as the 3SPN.2C DNA model [34]:

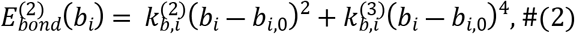

where *b_i_* and *b*_*i*,0_ have the same meaning as in Eq (1); 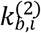 and 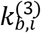 are the force constants in the quadratic and quartic terms, respectively.

### Angle potential

The harmonic angle potential is given by:

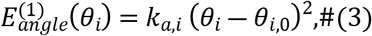

where *0_i_* is the bond angle formed by the adjacent three CG particles, *θ*_*i*,0_ is its reference value, and *k_a,i_* is the force constant.

The AICG2+ protein model [21] uses a knowledge-based angle potential, which is based on the Boltzmann inversion method [48],

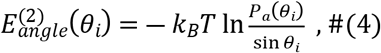

where *k_B_* is the Boltzmann constant, *T* is temperature in MD simulation, and *P_a_*(*θ_i_*) is an amino-acid type-dependent probability distribution of the angles analyzed from a set of PDB structures. 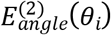 is further refined via the iterative Boltzmann algorithm. Practically, in MD simulations, we calculate the energies and forces using spline functions fitted to values in a table containing the energy and its derivatives at several points.

### Dihedral potential

The periodic proper dihedral potential is defined as follows:

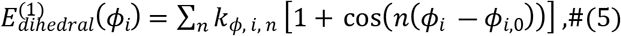

where *ϕ_i_* and *ϕ*_*i*,0_ are the dihedral angle and its reference value, respectively, *n* is an integer number that controls the periodicity of the function, and *k_ϕ,i,n_* is the force constant.

We also implement a Gaussian type dihedral potential, which is used in the AICG2+ [21] and the 3SPN.2C models [34]:

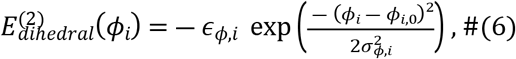

where *ϕ_i_* and *ϕ*_*i*,0_ have the same meaning as in Eq (5), *ϕ_ϕ,i_* controls the Gaussian width and *ϵ_ϕ,i_* is the force constant.

Similar to Eq (4), AICG2+ [21] also uses a knowledge-based dihedral term, which is based on the Boltzmann inversion potential [48]:

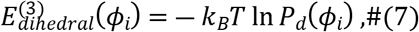

where *P_d_*(*ϕ_i_*) is the probability distribution of the dihedral angle. The 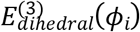 was further refined via the iterative Boltzmann algorithm. This is also calculated as a tabulated function for better performance. Notably, in each of the dihedral potentials described above, we introduced a recently proposed algorithm [49] to improve the robustness and numerical stability in the force evaluations.

### Nonbonded interactions

The structure-based Gō-like models for proteins usually use a 12-10 potential to model a native contact, defined as two residues containing heavy atoms at a distance less than a specific cutoff in the native structure [14,21]. The potential [14] is defined as:

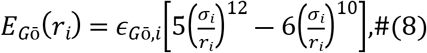

where is the distance between two CG particles in a native-contact pair, *σ_i_* is the native value of *r_i_*, and *ϵ_Gō,i_* is the force constant. Note that it is applicable not only to proteins but also to nucleic acids if their experimental structures are known.

The Lennard-Jones (LJ) potential (or 12-6 potential) is also widely used to describe pairwise nonlocal interactions and is commonly defined as:

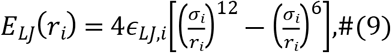
where *ϵ_LJ,i_* represents the minimum energy.

The LJ potential truncated at its energy minimum is a purely repulsive potential and can be used as the excluded volume interaction:

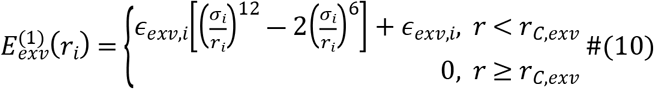

where *r_C,exv_* is the cutoff distance and has the same value as *σ_i_*. Note that the LJ forms in Eq (9) and Eq (10) are different in a coefficient of 2 for the 6th-power term. However, these two forms are mathematically equivalent when the parameters (*σ* and *ϵ*) are appropriately converted.

A simpler form of the excluded volume interaction is given by:

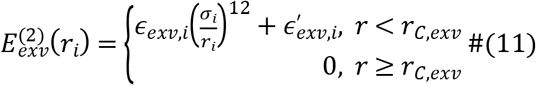

Other pairwise nonbonded interactions include the Gaussian potential:

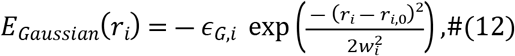

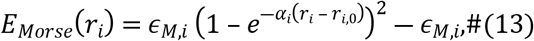

where *ϵ_G,i_* and *ϵ_M,i_* are the “depth”, and *w_i_* and *α_i_* are the “width” of the Gaussian and Morse potentials, respectively. In the 3SPN.2C model [34], the Morse potential is divided into two parts, the repulsive 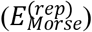 and the attractive 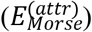, as defined in the following:

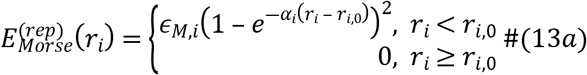

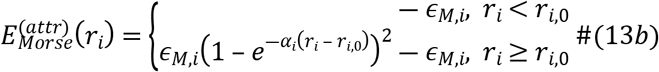

The electrostatic interactions are modeled using the Debye-Hückel theory:

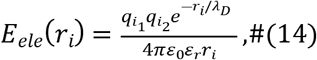

where *r_i_* is the distance between the charged particles *i*_1_ and *i*_2_, whose charges are *q*_*i*_1__ and *q*_*i*_2__, respectively. *ε*_0_ is the dielectric permittivity of vacuum. *λ_D_* is the Debye screening length. *ε_r_* is the relative permittivity of the solution, which is a function of the solution temperature *T* and salt molarity *C*: *ε_r_* = *e*(*T*)*a*(*C*), where *e*(*T*) = 249.4 – 0.788 *T* + 7.20 × 10^−4^*T*^2^ [50], and *a*(*C*) = 1 – 0.2551*C* + 5.151 × 10^−2^*C*^2^ −6.889 × 10^−3^*C*^3^ [51]. The Debye length is defined as: 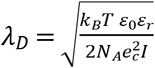, where *N_A_* is the Avogadro’s number, *e_c_* is the elementary charge, and *I* is the ionic strength of the solution.

### Modulating functions

When residue-level CG models are used to describe specific nonbonded interactions, such as the hydrogen bonds (HBs) or the π-π stacking, it is necessary to introduce multi-body potentials. For instance, in the 3SPN.2C model, the base-stacking, base-pairing, and cross-base-stacking interactions are described with the angle and torsion-angle dependent multi-body potential functions [23,34]. Specifically, the angle-dependent modulating function in the 3SPN.2C model is defined as [34]:

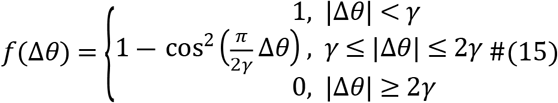

where Δ*θ* is the difference between an angle (*θ*) and its reference value, and *γ* controls the tuning range.

### Residue-level CG models

We next describe the CG models as the combinations of the basic interaction terms defined in the last section. We mainly focus on the energy function forms rather than the detailed parameters, which users can easily change in their MD simulations.

### The AICG2+ model for folded proteins

The energy function of the AICG2+ model [21] is defined as:

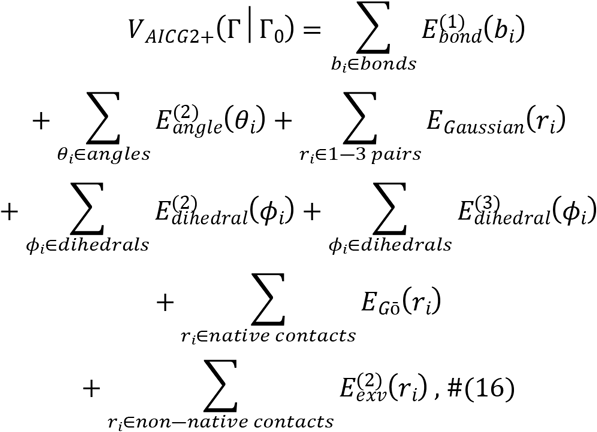

where Γ and Γ_0_ are the simulated and native structures, respectively. The reference values in each term are determined from the native values in Γ_0_, except for the non-native contact term. In the non-native contact interactions, *σ_i_* is dependent on the excluded volume radii of the particles in contact, using the combination rule of 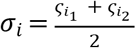, where *i*_1_ and *i*_2_ are the indices of the particles in the *i*-th non-native contact, with *ς*_*i*_1__ and *ς*_*i*_2__ as their residue type-dependent radii.

### The HPS and KH models for IDR

The two IDR models, HPS [28] and KH [29], share the same energy function [30]:

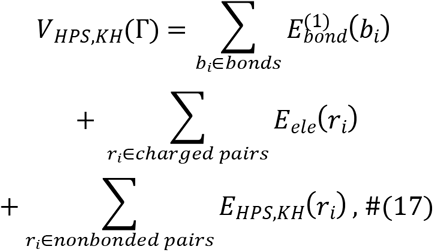

where Γ is the conformation of an IDP. *E_HPS,KH_*(*r_i_*) has the Ashbaugh-Hatch form [52]:

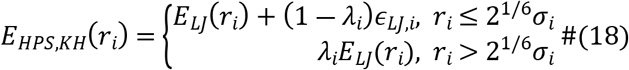

where *λ_i_* in the HPS model is the hydrophobicity scale, and in the KH model is a sign function to regulate the attractiveness or repulsiveness. All the other quantities, *r_i_*, *σ_i_*, and *ϵ_LJ,i_*, have the same meaning as defined in Eq (9). Specifically, *ϵ_LJ,i_* and *λ_i_* are based on the amino acid hydrophobicity in HPS [28] or the Miyazawa-Jerningan potential in KH [29,53], respectively.

The same potential functions are also used in the HPS RNA model [32], which has been developed to study the co-condensation of RNA and protein IDRs. In the HPS RNA model [32], the resolution is only one bead for one nucleotide, which is lower than the other nucleic acid CG models in GENESIS. Accordingly, the bond lengths and particle radii are also larger than the other models. Note that GENESIS also provides a structure-based three-bead-per-nucleotide RNA model, as described in a later section.

### The 3SPN.2C model for DNA

The potential energy function of the 3SPN.2C model [23,34] is defined as follows:

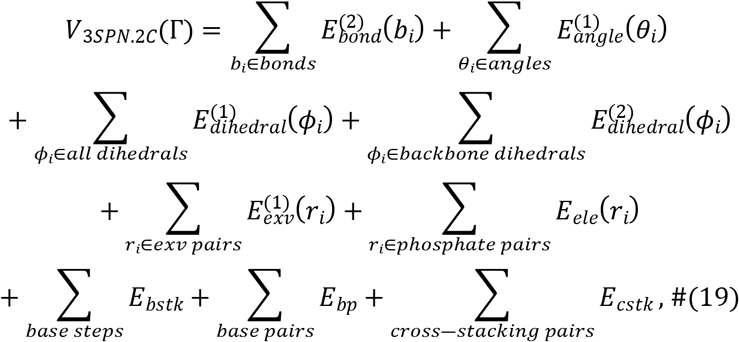

where Γ is the conformation of the DNA molecule, and “exv pairs” are the nonbonded particle pairs that are not involved in either base-pairing or stacking interactions. The three terms for base-base interactions, *E_bstk_, E_bp_*, and *E_cstk_*, are defined as multi-body energy functions, using the basic equations described in the previous section (Eq (13a), (13b) and (15)):

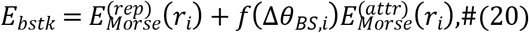

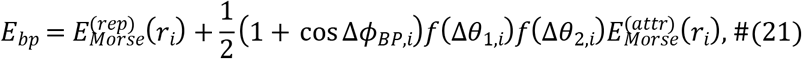

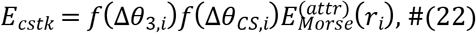

where *r_i_* represents the distance between the two interacting bases, and the angles (*θ_BS,i_*, *θ*_1,*i*_, *θ*_2,*i*_, *θ*_3,*i*_, and *θ_CS,i_*) and dihedral angles (*ϕ_BP,i_*) are formed by the surrounding sugar and phosphate sites (for details of the definition, please refer to [34]).

### The structure-based model for RNA

In addition to the one-bead (per-nucleotide) HPS model, we also implement a structure-based three-bead model for RNA [22], whose parameters were determined with the fluctuation matching method [54]. The energy function of this model [22] is defined as:

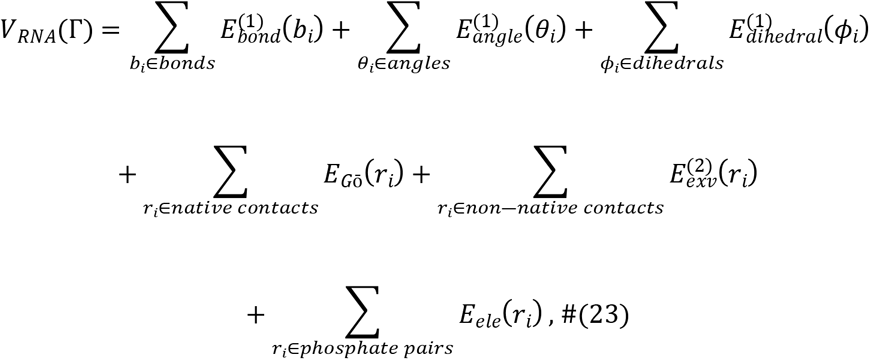

where Γ is the conformation of the RNA molecule. The electrostatic term, *E_ele_*, is optional in the intra-RNA interactions but required for protein-RNA interactions [22]. In the later case, *r_i_* in *E_ele_*(*r_i_*) represents the distance between a charged amino-acid residue and an RNA phosphate.

### The PWMcos model for protein-DNA interaction

The protein-DNA binding is generally considered in two parts: the sequence-nonspecific interactions between amino acids with the DNA backbone groups (mainly the electrostatic interactions) and the sequence-specific interactions between amino acids and DNA bases. The PWMcos model can describe the latter, incorporating the PWM information into the structure-based interactions [27]. The model first defines a list of DNA-binding protein residues (DB-*C_α_*s) forming contact with DNA in the native structure, and then the potential energy is then given by [27]:

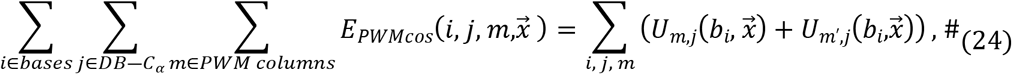

where *m*′ is the base in the complementary strand that forms base-pair with base *m, b_i_* is the base type of any base *i* (*b_i_* ∈ [*A, C, G, T*]), and 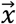 is the coordinates of particles in each conformation. The 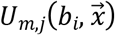 function is defined as:

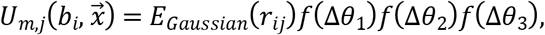

where *r_ij_* is the distance between the *i*-th and the *j*-th *C_α_*, and *θ*_1_, *θ*_2_, and *θ*_3_ are three angles defined by the surrounding particles (for details, please refer to [27]). *E_Gaussian_*(*r*) and *f*(Δ*θ*) functions are defined in Eq (12) and (15), respectively. The energy coefficient of *E_Gaussian_*(*r*) (*ϵ_G,i_* in Eq (12)) is defined as a function of base type *b_i_* based on the PWM:

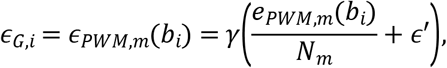

where *e_PWM,m_*(*b*) is the element in the mth column of the PWM and in the row corresponding to base type *b, N_m_* is the total number of protein contacts formed with the *m*th base pair, *ε*′ is a hyperparameter that shifts the absolute value of the function, and *γ* is a scaling factor to change the strength of the interaction. Notably, a variation of the PWMcos has been used to model sequence-non-specific hydrogen bonds between protein and DNA backbone [25]:

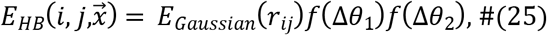

where *r_ij_* represents the distance from the *i*-th phosphate (instead of base) to the *j*-th *C_α_*, and *θ*_1_ and *θ*_2_ are the angles defined by the local particles around the target phosphate and *C_α_* atom in the configuration with coordinates 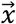.

### Summary of the potential energy functions available in GENESIS

Although the models mentioned above are developed independently, some have been proved to work well with each other. For instance, the association of the AICG2+ and the 3SPN.2C models has shown the capability to study the protein-DNA binding systems [24,26]. Between proteins and DNA, electrostatic interactions contribute most to the sequence-nonspecific affinity. For the accuracy of electrostatics modeling, the RESPAC method was developed to calculate the surface charge distribution of proteins [55]. Furthermore, the angle-dependent HB potential enables modeling the rotation-coupled and uncoupled sliding of nucleosomal DNA [25,56]. The PWMcos method also facilitates the target search of transcription factors on both linear DNA and nucleosomes [27,57].

In addition to these well-developed methods, GENESIS also allows the combinations of different models. In Table 1, we list all the available CG models in GENESIS 1.7.0 in mixtures of proteins, DNAs, and RNAs.

**Table 1.**
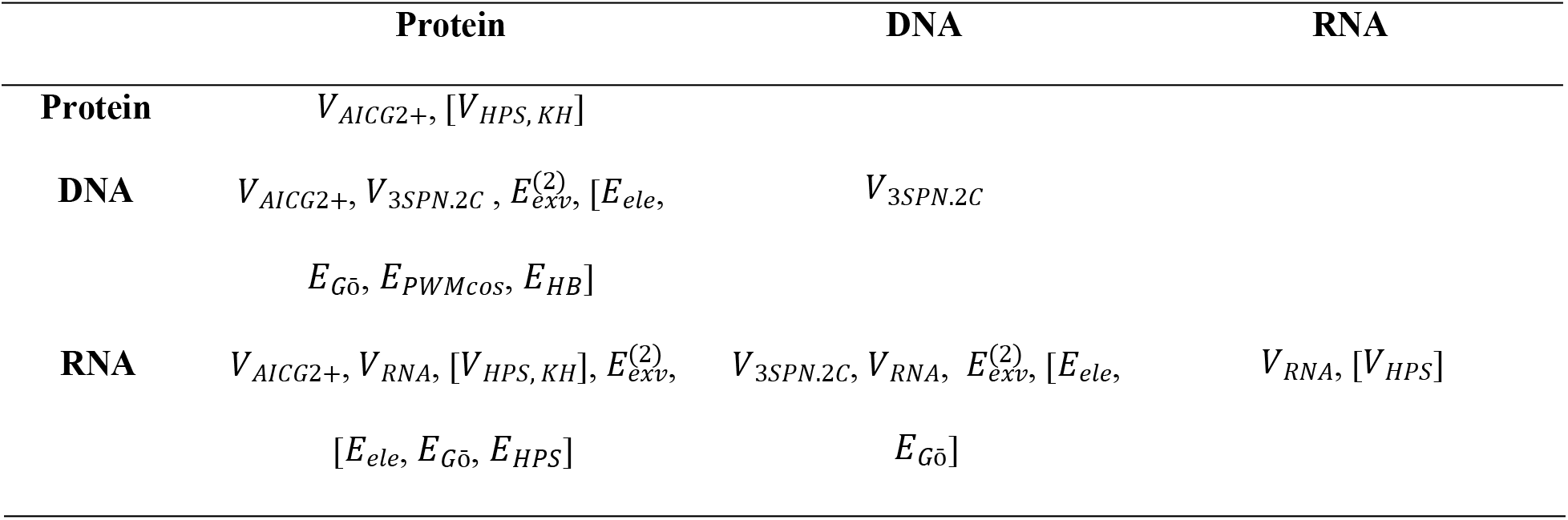
Available CG models in heterogeneous biomolecular systems.

The interactions written in brackets are optional.

### Information flow

Fig 2 shows a flowchart for running CG simulations in GENESIS. We roughly divide the whole procedure into three steps: (i) preparation, (ii) simulation, and (iii) analysis. GENESIS provides tools to achieve the task in each step. In the preparation stage, we use a collection of scripts, “GENESIS-CG-tool”, to translate experimental results into CG topology and coordinate files. Together with the MD control file, these files are then parsed by one of the GENESIS MD engines, atdyn, to perform energy/force calculations and time integrations. As the output data from MD simulations, GENESIS produces a log file of system properties including temperature, system size, and potential energy. The output coordinates from MD simulation are written in a trajectory file with the DCD format, which are processed by the GENESIS analysis tools and can be visualized by VMD [58], PyMOL [59], or other molecular graphics software. Here, we mainly focus on our developments in the first two stages, since the analysis using the DCD format files and analysis tools is common to those of all-atom MD simulations.

**Fig 2.**
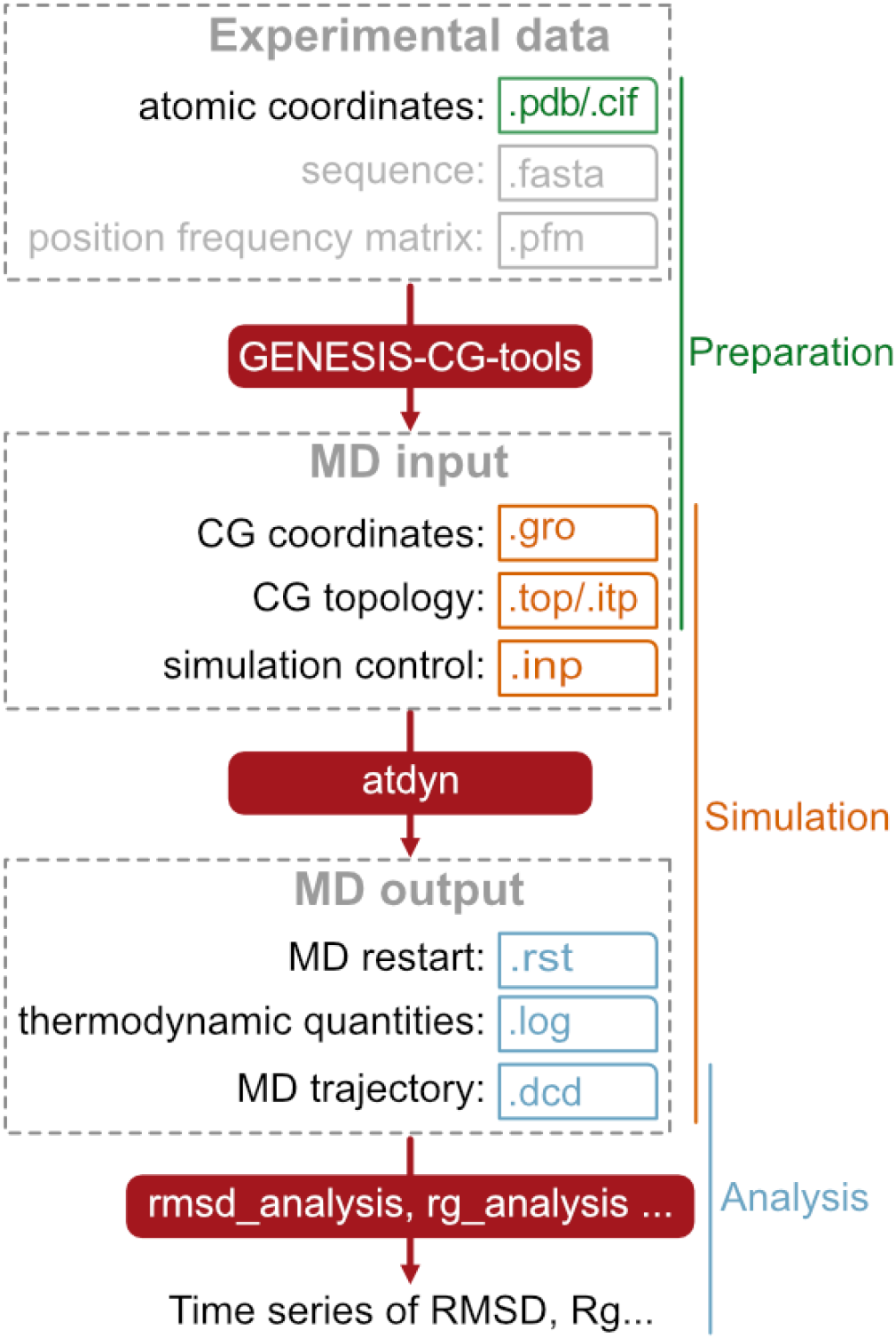
Information flow in GENESIS CG MD simulations. Dashed boxes represent collections of files used together as input or output at a particular step. Solid red boxes are used for the executable tools provided by GENESIS. Small boxes include typical filename extensions.

### Preparation

We next explain the usage of the GENESIS-CG-tool, which can read structural information and generate force field parameters and coordinates for MD simulations. As input, the structural information is in the PDB format, which includes atom coordinates determined by experiments such as X-ray crystallography, NMR, or Cryo-EM. As output, we decided to use a unified GROMACS-like file format for all the CG models. Particularly, the output of GENESIS-CG-tool includes a major topology file (.top) and a bundle of molecule-specific files (.itp), as well as a coordinate file for all the particles in the system (.gro) (Fig 2). The molecule-specific topology (.itp) files contain the information of intra-molecular interactions, such as bond, angle, dihedral, and Gō-type native-contact interactions. Each term includes the indices of the involved particles, reference values of the variables, and the force parameters. Additionally, an integer variable (called the “function type”) is used to distinguish the various potential functions of the same interaction type. For instance, we set the function types of 1 and 21 to 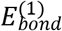 (Eq (1)) and 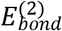 (Eq (2)), respectively.

GENESIS-CG-tool was developed with Julia [47], a fast, open-sourced, and reproducible programming language. The project’s main entrance is a script called “aa_2_cg.jl”, which works in the command-line environment of UNIX-like shells. The script takes a PDB file name as the positional argument and several model-related optional arguments. During the processing, the atomistic coordinates are read from PDB files. Then, local and nonlocal interactions, including their native values, are detected from the PDB structure. Other parameters, such as force constants, are determined by the models specified for each molecule.

As a simple example, assuming that one wants to prepare CG MD input files for a protein using a PDB file, “PRO1.pdb”, the command “aa_2_cg.jl PRO1.pdb” will generate “PRO1_cg.top” and “PRO1_cg.itp” topology files and “PRO1_cg.gro” coordinate file (Fig 2 and S1 Fig). The AICG2+ model is used for proteins by default, which can be changed with the option “--force-field-protein”.

The 3SPN.2C model of DNA uses reference structures that include sequence-dependent geometric features. These structures can be generated by software such as 3DNA [60]. In GENESIS-CG-tool, with no requirement of external packages, we provide a simple “sequence-to-structure” script, which reads in DNA sequences and directly outputs both atomistic PDB and CG topology/coordinate files for dsDNA, including the necessary sequence-dependent information (see S1 Fig).

For heterogeneous systems that contain more than one type of biomolecules, the molecule-specific “.itp” files can be linked into the “.top” file, using the “#include” keyword.

Notably, recent improvements in experimental techniques such as cryo-EM have fostered more high-resolution structures of the huge biomolecule complexes, which are valuable resources in MD simulations. The spatial scales of these structures, however, outstrip the capacity of the traditional PDB file format. Another format, PDBx/mmCIF [61], has been used to store coordinate information of large structures. GENESIS-CG-tool also accepts files of the PDBx/mmCIF format as input. Besides, we also noticed that detecting the Gō-like native contacts is rather time-consuming for large protein complexes. We utilize the Julia thread parallelization to accelerate this procedure in the GENESIS-CG-tool.

We have packaged GENESIS-CG-tool as a part of GENESIS v1.7.0. We have also deployed the GENESIS-CG-tool in an individual Github repository: https://github.com/noinil/genesis_cg_tools. Users can refer to the online wiki page of this project for more details on the usage and available file formats.

### MD algorithms

In GENESIS v1.7.0, there are two MD programs: spdyn and atdyn. Spdyn is parallelized using the mid-point cell method [62], one of the domain-decomposition schemes; while atdyn is parallelized with the atomic decomposition scheme [45,46]. We implement the residue-level CG models only in atdyn because of the feasibility of implementations and the number of CG particles required in most CG MD simulations. In atdyn, the Clementi Gō [14], Karanicolas-Brooks Gō [63], Domain-enhanced Model (DoME) [64], and dual/multi-basin Gō models [65] are also available and use the same topology and coordinate file formats as described in the previous section. Since atdyn is already a well-developed framework, the original source code is reused as much as possible to implement the residue-level CG models. In the following, we describe the newly introduced functions and optimization schemes only.

### Cutoff scheme of the nonbonded interactions

The nonbonded interaction terms are the most time-consuming computations both in all-atom and CG MD simulations. Cutoff schemes are used for the nonbonded terms in CG MD simulations to reduce the computation time. For relatively short-range nonbonded interactions, such as 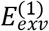 and 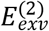, cutoff values are already included in the definition (see Eq (10) and (11)). In contrast, the other interactions do not have generally established cutoff values. We provide three parameters to control the cutoff lengths of the *E_bp_*, *E_LJ_*, and *E_ele_* terms in our implementations. We set the default cutoff value of *E_bp_* to 18Å, following the original 3SPN.2C model [23,34]. We use a value of 52Å for *E_ele_*, which assures that the absolute value of the energy at the cutoff distance to be smaller than 10^−4^ kcal/mol (calculated at the temperature of 300K and with an ionic concentration of 150mM, for two charges of ± 1 *e^−^*). This cutoff value of *E_ele_* is slightly larger than the one used in the original 3SPN.2C model and guarantees accuracy [34]. As for the LJ potential used in the HPS/KH IDR models, the parameters *ϵ_LJ,i_* and *σ_i_* are dependent on residue-types [30]. Therefore, we determine the LJ cutoff using a strategy that, for the i-th pair, the absolute value of the energy at the cutoff distance is as small as 10^−4^ times of the energy minimum: *E_LJ_*(*r_C,LJ_*) = 10^−4^*ϵ_LJ,i_*. This gives a value of *r_C,LJ_* – 5.849*σ_i_*. We then choose the largest *E_LJ_* from the HPS/KH models to determine a unified value of 39Å as the cutoff for *E_LJ_* Note that these default values are considered as the “safest” ones. The users may be able to change them to smaller values by carefully evaluating the balance between computational efficiency and accuracy.

Several nonbonded interaction terms are considered only for pre-defined pairs of CG particles, such as the DNA base-stacking (*E_bstk_*) and the Gō-type native contacts (*E_Gō_*). For these terms, we don’t apply the cutoffs but directly calculate all the interactions.

### Neighbor lists

Neighbor lists are used to maintain a list of particles that can participate in nonbonded energy/force interactions within a few MD integration steps. We set the neighbor list distances (*r_P_*, also called “pair-list distance” in GENESIS) to be roughly 5Å larger than the largest cutoff distance (*r_C_*). A complete list of the default cutoff and neighbor list distances can be seen in Table 2.

**Table 2.**
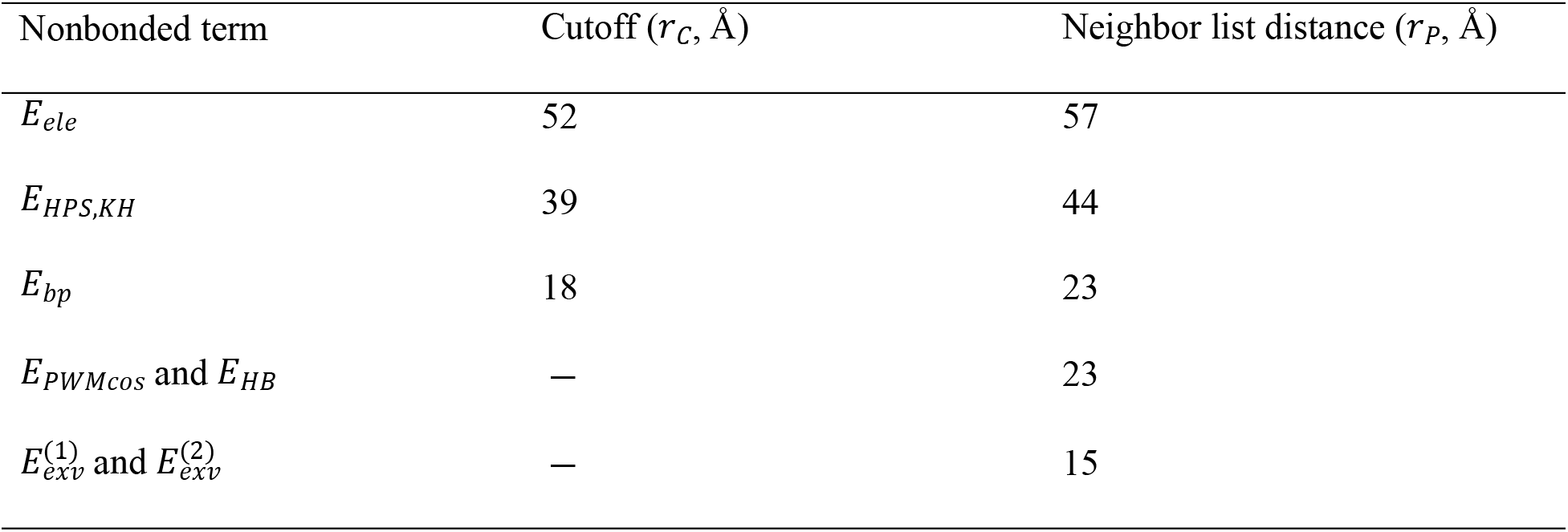
Default values for the nonbonded cutoff and pair-list distances.

The cutoff of *E_PWMcos_* and *E_HB_* are given by *r*_*i*,0_ +5Å (*r*_*i*,0_ is defined in Eq (12)). The cutoff of 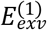 and 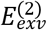 are defined in Eq (10) and (11), respectively.

We utilize the cell linked list strategy to construct the neighbor list. Fig 3 illustrates how we determine the neighbor lists of different nonbonded interaction terms. During every update of the neighbor lists, the system is first divided into small cells. In each cell *C_i_* and for each interaction term, we construct a “particle list”, *L_nb–term_*(*C_i_*), which contains the particles located in *C_i_* and that are involved in the interaction (“nb-term”). For example, *L_ele_*(*C_i_*) is a list of all the charged particles in the cell *C_i_*. We then determine the “neighboring cells” for each cell: 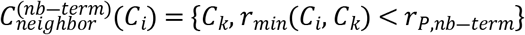, where *r_min_*(*C_i_, C_k_*) is the minimum distance between the cells *C_i_* and *C_k_*. As shown in Fig 3A, relatively short-range interaction terms such as 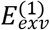 or 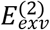 have a small number of the neighboring cells, whereas those with longer cutoff distances, such as *E_ele_*, will have more neighboring cells.

**Fig 3.**
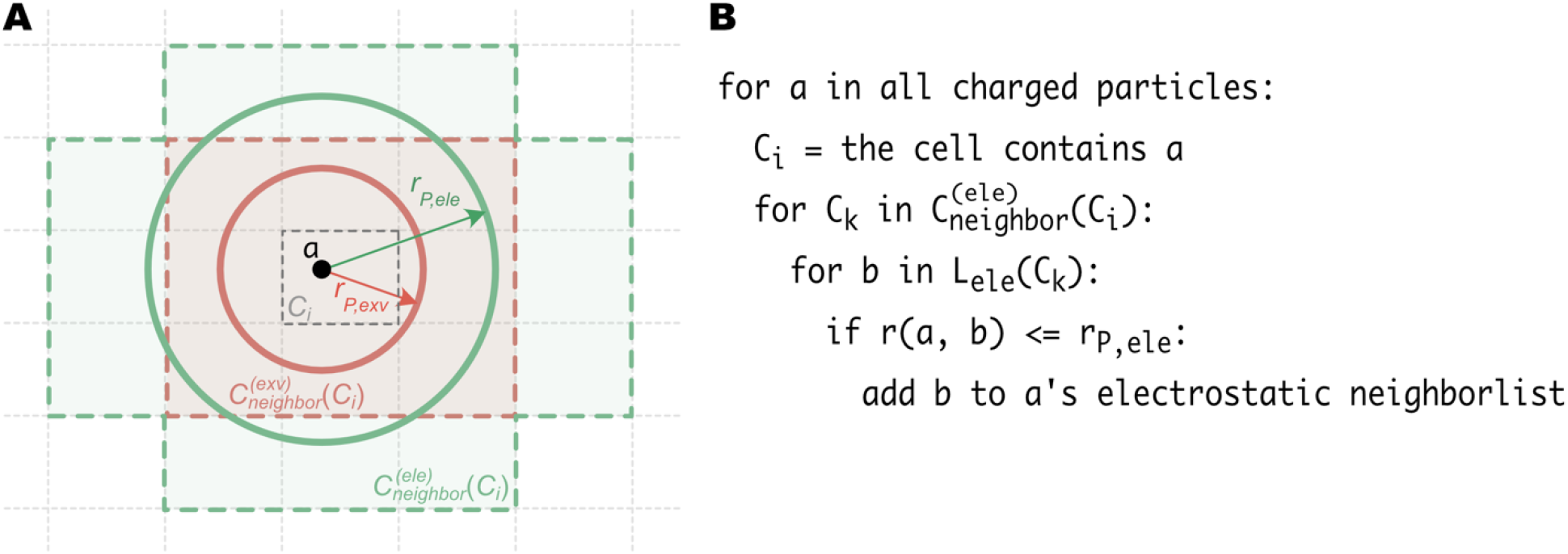
Using the cell linked list method to construct multiple neighbor lists. (A) An example of determining the neighbor list for *E_exv_* (representing either 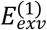 or 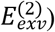) and *E_ele_* on a two-dimensional space. The gray dotted lines show the edges of the cells. The center cell with dark dashed edges (*C_i_*) contains the particle *a* (black dot) whose neighbor list is to be determined. *r_P,exv_* and *r_P,ele_* are the neighbor-list distances for *E_exv_* and *E_ele_,* respectively. The neighboring cells for the *E_exv_* term 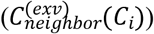 are represented by the red cells and surrounded by the red dashed lines. Those for the 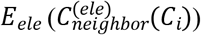 are shown as the green area edged by green dashed lines. The red and green circles indicate the range of final neighbor lists for *E_exv_* and *E_ele_*, respectively. (B) Pseudo code of the algorithm to determine a neighbor list of the electrostatic interaction. *L_ele_*(*C_k_*) is a list of all the charged particles in cell *C_k_*.

After defining the “particle list” (*L_nb–term_*(*C_i_*)) and “neighboring cells” 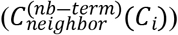 of each cell, the neighbor list for each particle can be determined by the algorithm in Fig. 3B (for convenience, here we use electrostatic interaction as an example). The neighbor lists are updated every 20 steps in the CG simulations. Using individual particle list and neighboring cell for each nonbonded interaction term, we can minimize the pairwise distance calculations. First, as described above, we can avoid do loops over unnecessary cells for short-ranged interactions by using the potential-dependent neighboring cells. Besides, in each cell, we just consider a pre-assigned subgroup of particles instead of all the particles.

### Time Integration

GENESIS atdyn has provided various integrators and thermostats/barostats. For the current CG implementation, we employ the velocity-Verlet integrator coupled with the Langevin thermostat. In the velocity-Verlet algorithm, coordinates (**r**_*i*_) and velocities (**v**_*i*_) are updated following:

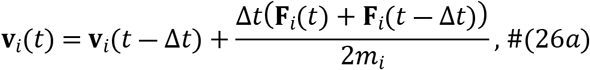

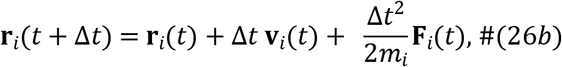

where *t* is the simulation time, Δ*t* is the integration step size, *m_i_*, **r**_*i*_, **F**_*i*_, and **v**_*i*_ are the mass, coordinate, force, and velocity of the *i*-th particle, respectively. In the Langevin thermostat, the force **F**_*i*_ is calculated by:

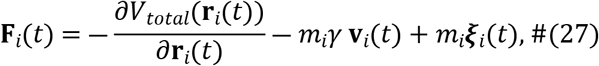

where *V_total_* is the total potential energy and *γ* is the friction constant. ***ι***(*t*) is the Gaussian noise vector satisfying:

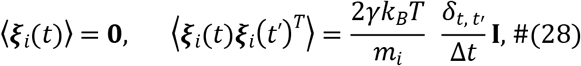

where *k_B_* is the Boltzmann constant, *T* is temperature, *δ_t,t′_* = 1 when *t* = *t*′ and 0 otherwise, and **I** is an identity matrix of size 3 × 3. ***ξ**_i_*(*t*′)^*T*^ is the transpose of ***ξ**_i_*(*t*′) and the product follows matrix multiplication.

We use 10 femtosecond (fs) as the literal time step size (Δ*t* in Eq. 26), which is determined by the highest vibration frequency of the particles. On the other side, the time scale mapping of large-scale conformational dynamics of biomolecules is system-dependent and requires a careful comparison between simulated and experimental physical quantities [17].

### Performance Optimization

Next, we optimize the performance of CG MD simulations implemented in GENESIS. At first, we optimize the nonbonded interactions, which are the most time-consuming parts in both residue-level CG and all-atom force-field models. There are two common strategies in the optimization. The first one is the vectorization of the most inner loops for utilizing as many SIMD (single instruction and multiple data) instructions as possible. The second one is to minimize the number of computations by using conditional statements. In contrast to all-atom models, using “conditional” statements is not avoidable due to the complex potential energy function forms. In our experience, the ratio *r*_*P*_^3^: *r*_*C*_^3^ gives us a good criteria whether the calculation would be vectorized or not using “conditional” statements. For instance, in the case of *E_ele_,* the ratio is about 1.3, and it is better to apply vectorizations to the inner loop without any “conditional” statements. The ratio in 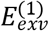 or 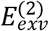 evaluations are greater than 3.5, and it is better to optimize by minimizing the calculation amount with conditional statements. The ratio in *E_LJ_* or *E_bp_* is less than 3, but we use conditional statements because the energy expression form itself is changed in HPS, KH, or 3SPN.2C models.

Another computational bottleneck in CG MD simulations is the generation of random numbers with Gaussian distributions when using the Langevin thermostat. This is mainly because of the replicated data MPI parallelization scheme used in GENESIS atdyn, in which all processes have a copy of the coordinate data. By keeping the replicated data scheme, we assign MPI parallelization in random number generation and apply “MPI_Allgatherv” to collect the information. We do not apply MPI parallelization to integrate coordinates or momenta because the communication time is comparable or even more time-consuming than the integration itself. Instead, OpenMP parallelization is applied in integrations.

## Results

### Benchmark of MD simulations with residue-level CG models

We first examine the usage of memory and the performance benchmark in MD simulations with residue-level CG models. The memory usage was examined with a single process and a single thread on a machine with 93GiB RAM (random access memory). The CPU benchmarks were executed with 5 OpenMP threads and various MPI processes on a computer server with Intel Xeon Gold-6148 (2.4GHz) CPU (20 cores per CPU and two CPUs per node) and InfiniBand EDR networking. GENESIS was compiled with the Intel compiler in couple with Intel MPI version 2019.5.281. Fig 4 shows the results of the memory benchmark (Fig 4A) and CPU benchmark using different biological systems (Fig 4B to 4D).

**Fig 4.**
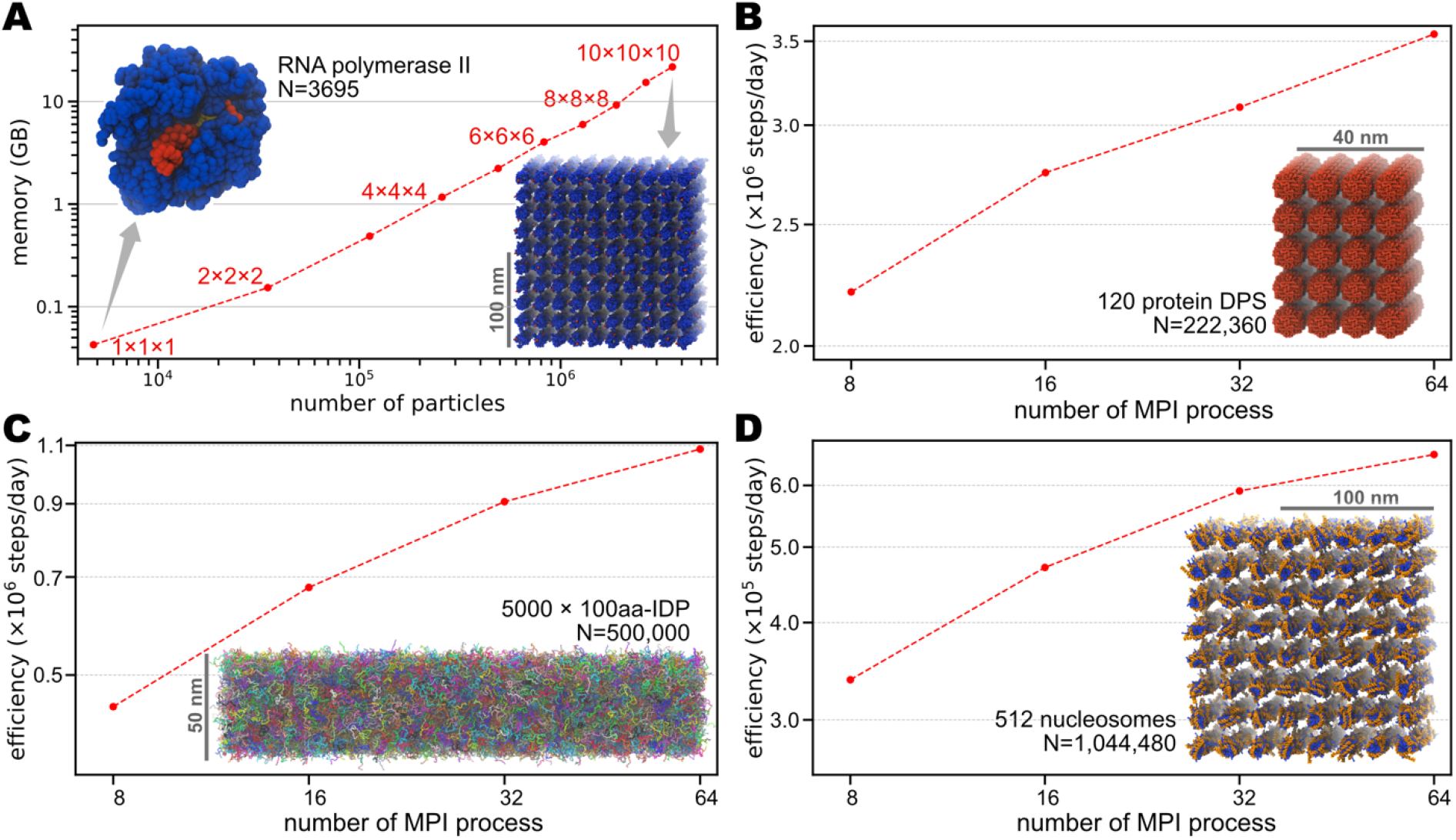
Benchmark of the GENESIS CG simulations. (A) Memory benchmark of the RNA Polymerase II (PolII) systems. A single PolII has 3,695 CG particles, which is then duplicated into a *n* × *n* × *n* (*n* = 1,…,10) grid space. The insets show the structure of a single PolII (left) and 1000 PolIIs (right). Protein is shown in blue, DNA in red, and RNA in yellow. (B) CPU benchmark of a system containing 120 DPS proteins. The total number of CG particles is 222,360. (C) CPU benchmark for a system of 5000 chains of 100aa IDPs. The total number of CG particles is 500,000. (D) CPU benchmark for a system including 512 nucleosomes. The total number of CG particles is 1,044,480. The insets of (B), (C), and (D) show the initial structure of each system, respectively. N represents the total number of CG particles in each system. For all the CPU benchmarks, we used 5 OpenMP threads.

To test the memory consumption, we built systems consisting of different numbers of RNA Polymerase II (PolII). The smallest system has only one PolII and contains 3695 CG particles (3542 from protein, 124 from DNA, and 29 from RNA), based on the X-ray crystallography structure (PDB ID: 1R9T) [66]. We then duplicated the single PolII to create systems of different sizes, ranging from thousands to millions of particles. In all these tested systems, we used *V_AICG2+_* for proteins, *V_3SPN.2C_* for DNAs, and *V_RNA_* for RNAs. For proteins, we use integer charges distributed on the charged residues ( *+e* for Arg and Lys, *−e* for Asp and Glu). For the interactions between different biomolecule types, we applied 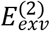 and *E_ele_* as general nonspecific interactions, and *E_Gō_* for native contacts between protein-RNA, protein-DNA, and RNA-DNA to maintain the whole structure. All the cutoff and pair-list distances are set to the default values (see Table 2). As shown in Fig 4A, we carried out simulations with *n*^3^ (*n* = 1,…10) duplicated PolIIs to track the memory usage. The consumed memory scales with the system sizes, from ~40 MB for a single PolII to ~22 GB for 1000 PolIIs. Notably, on an ordinary computer with 10GB memory, we can run GENESIS CG MD simulations for a system with about 2 million particles.

To examine the strong scaling of GENESIS CG simulations, we prepare three systems comprised of different types of biomolecules:

1. 120 copies of the protein DPSs [67]. In total, 222,360 CG particles modeled with *V_AIGG2+_*;
2. 5000 chains of IDPs, each consisting of 100 amino acids. In total, 500,000 CG particles modeled with *V_HPS_*;
3. 512 copies of nucleosomes [68]. In total, 1,044,480 particles, using *V_AICG2+_* for proteins, *V_3SPN.2C_* for DNAs, and 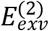, *E_ele_* for protein-DNA interactions.

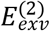 and *E_ele_* are used as general non-specific interactions for systems 1) and 3); and *E_HB_* is additionally applied between protein and DNA in system 3). For all the cutoff and pair-list distances, we use the default values listed in Table 2. The benchmark results for these three systems are shown in Fig 4B-D, respectively. The results suggest that at least up to 64 MPI processes (8 nodes), the parallel performances are scalable in all three systems. Although this does not show perfect scalability, it is useful for simulating large biological systems. We discuss possible reasons for the insufficient scalabilities of our implementation and future plans to improve the performance in the Discussion section.

### Application 1: protein target search on DNA

The AICG2+ protein model (*V_AICG2+_*), and the 3SPN.2C DNA model (*V_3SPN.2C_*), in combination with the sequence-specific (*E_PWMcos_*) or nonspecific (*E_HB_*) protein-DNA interactions, can be used to study the binding, diffusion, and conformational changes of protein-DNA complexes [24,25,56,57]. As a simple example, we performed MD simulations of the sex-determining region Y protein (sry) [69] and its target DNA. The modeling and simulation results are shown in Fig 5.

**Fig 5.**
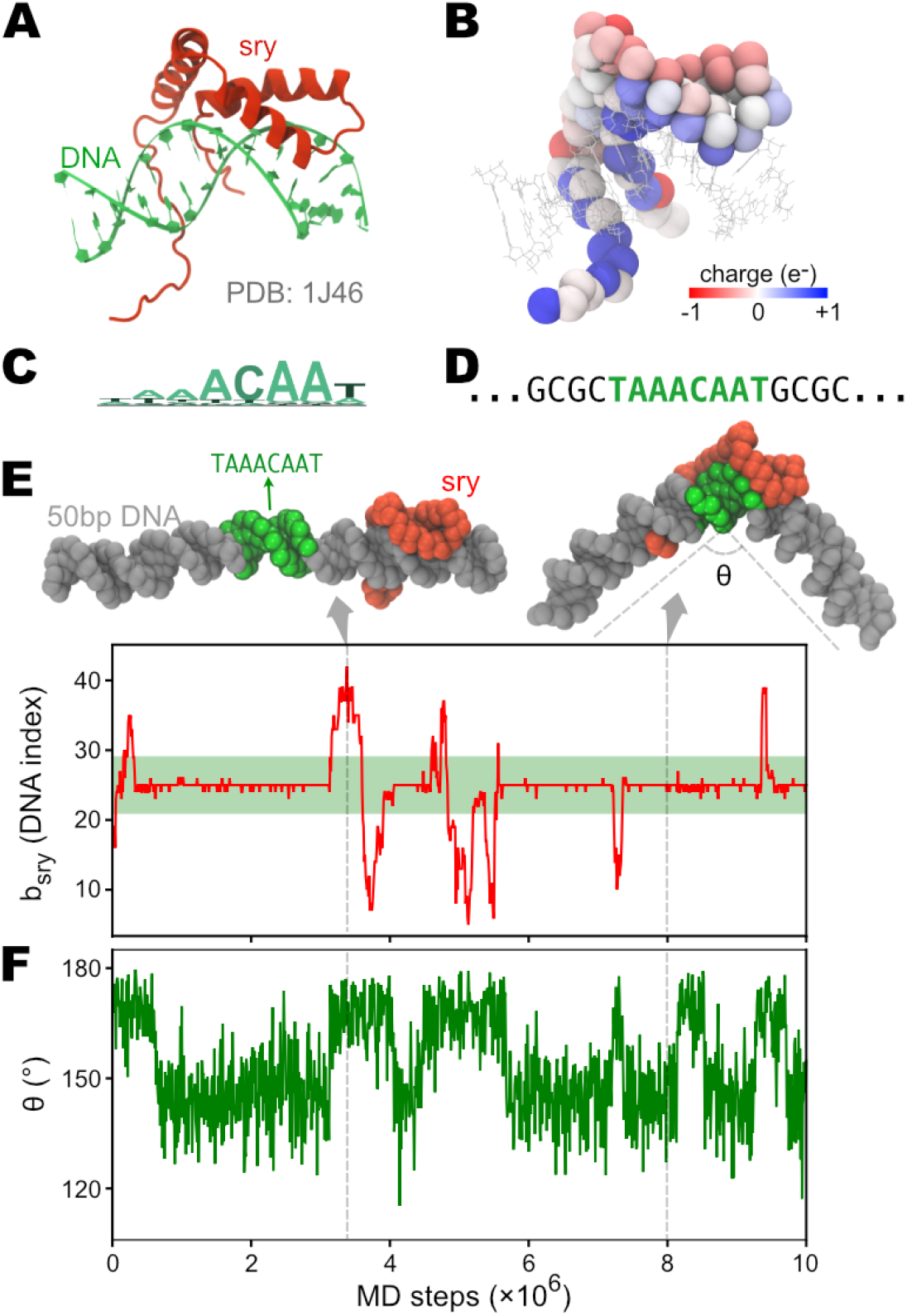
CG MD simulations of the DNA target search process by protein sry. (A) Solution NMR structure of sry binding on DNA (PDB entry: 1J46). (B) Coarse-grained sry with the C_α_ particles colored by the partial charges (determined using the RESPAC method). DNA structure from the PDB is also shown as lines for reference. (C) Sequence logo of sry’s DNA target. (D) Sequence of DNA used in the CG simulations. The consensus sequence “TAAACAAT” is inserted at the center of a 50bp poly-CG DNA. (E) Time series of sry’s binding position on DNA (*bsry*). The green region represents the consensus sequence. (F) Time series of the bending angle of DNA (θ). Two representative structures of sry binding at the poly-CG region or the consensus region are shown on top of (E). In these structures, DNA is colored in gray, except for the consensus sequence region in green. Sry is shown in red.

Sry uses a conserved High-Mobility Group (HMG) domain to recognize its consensus DNA sequence, “TAAACAAT” [70]. The HMG domain intercalates into the DNA minor groove and forms contacts with the bases. This type of minor groove binding of sry results in a sharp bending of the DNA [71], which plays a crucial role in regulating the transcription of its target gene [72]. Starting from the solution NMR structure of the sry-DNA complex (PDB entry 1J46 [71], see Fig 5A), we generated the CG model of sry using the GENESIS-CG-tool. We used the AICG2+ model (*V_AICG2+_*) for the whole protein. Specifically, we applied Gō-type native contact interactions (*E_Gō_*) to the folded HMG domain (residue 9-70) and assigned only local flexible potentials (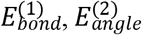, and 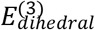) to the N-terminal (residue 1-8) and C-terminal tails (residue 71-85). We also employed the RESPAC method [55] to calculate the partial charge distribution on the surface residues of sry (see Fig 5B and S2 Fig). As for the DNA, we designed a 50bp poly-CG sequence, with the 8-bp piece “TAAACAAT” inserted at the center (Fig 5C and 5D). The atomistic structure of the dsDNA was then constructed from this 50-bp sequence using the 3DNA package [60]. The 3SPN.2C CG topology and coordinate files were then generated from the atomistic structure, using the GENESIS-CG-tool. For sry to find its consensus sequence in the simulations, we utilize the PWMcos model (*E_PWMcos_*) to incorporate information from the position frequency matrix (PFM, downloaded from JASPAR database [73] profile MA0084.1 [70]). For details of the model parameters, please see Supplementary Methods.

After getting the input files, we carried out MD simulations of the sry-DNA complex for 10^7^ steps. In the initial structure, sry was placed around 70Å away from the DNA. Soon after the simulation started, sry was bound onto DNA and began the sliding. We monitored the binding site of sry on DNA (Fig 5E) and the bending of DNA (Fig 5F). As can be seen, sry is preferentially bound to the consensus sequence (represented by the green region in Fig 5E) and occasionally left this target with a quick diffusion on the DNA. As expected, we observed coupling between the sequence-specific recognition and DNA bending caused by the intercalation of the HMG domain. These results show that the CG models (AICG2+, 3SPN.2C, and PWMcos) for protein-DNA interactions have been correctly implemented in GENESIS and can be used to study similar biological systems.

### Application 2: phase behaviors of IDR and RNA

The HPS and KH IDR models have been used to study the phase behavior of IDRs [30]. Here, we used the IDR from protein Fused in Sarcoma (FUS) [74] to show that our implementation of these models in GENESIS can simulate the condensation of proteins. We first simulated a single chain of the FUS IDR (163aa, see Supplementary Methods for the sequence). The final structure of the single-chain simulation was then duplicated to create a 120-chains system. We carefully arranged the multiple chains so that the system had a boundary size of 18*nm* × 18*nm* × 200*nm*, which is for the slab sampling method. We performed a 10^7^-step simulation, as shown in Fig 6A and 6B. During the simulation, we monitored the density change of the CG particles along the *z*-axis (the longest dimension). As shown in the top panel of Fig 6A, at the beginning of the simulation, the FUS IDRs formed relatively small and low-density clusters. As simulation time increased, we observed several merge events of the condensates. After ~ 3.5 × 10^6^ steps, all the FUS IDRs went into the same cluster, and the density finally reached around 5 particles per *n m*^3^ (Fig 6A and 6B). This simple example shows the possibility of using the HPS model in GENESIS to explore the physical mechanisms of IDR condensation. Besides, we noticed that there are some recent updates of the parameters of the HPS model [31,75]. These changes of parameters can be easily used with GENESIS by modifying the parameter files, with no need to touch the code. In this sense, GENESIS can also serve as a handy framework for the users to re-calibrate their own set of parameters.

**Fig 6.**
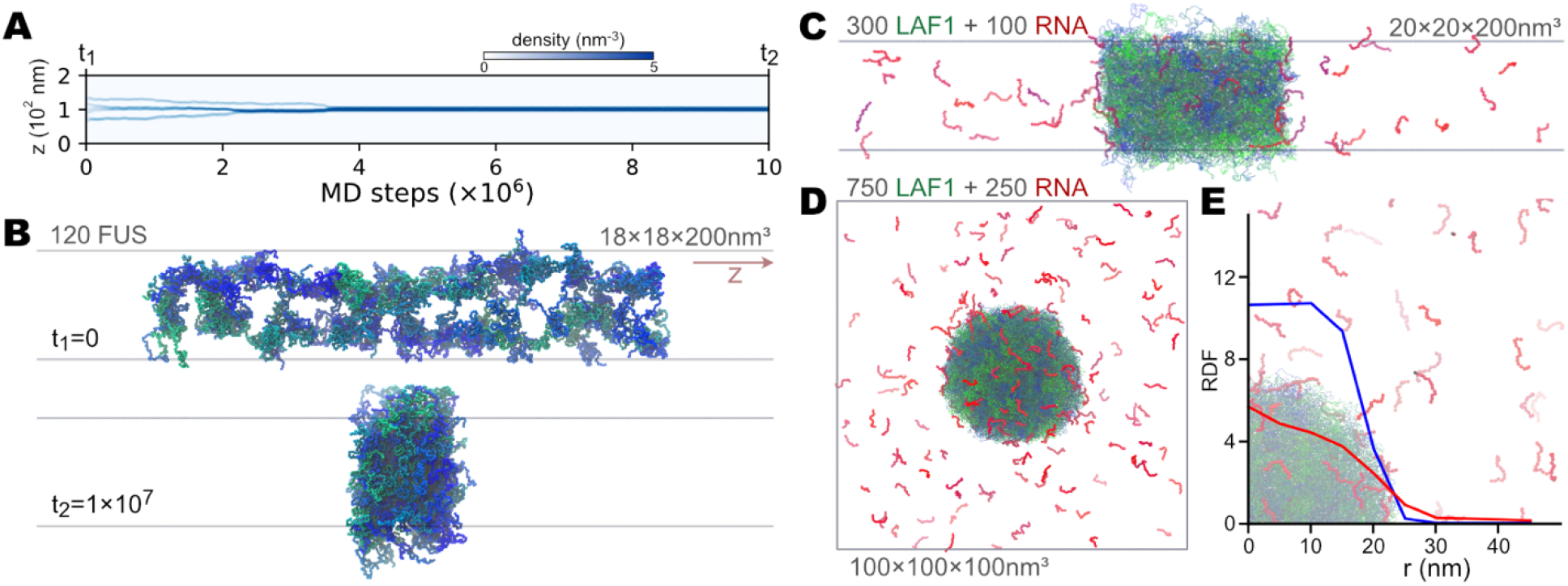
CG MD simulations of the condensation of IDRs and RNAs. (A) and (B) show the slab simulation results of 120 FUS IDRs. (A) Time series of FUS density along the long axis (*z*). The intensity of the blue color represents the density, as shown in the color bar above. (B) The first and last structure of the MD trajectory shown in (A). The FUS IDRs are shown as tubes. Different colors from green to blue are used to distinguish chains. (C), (D), and (E) show the simulation of the mixture of LAF1 IDRs and 15nt poly-A (A15) RNAs. (C) The last structure of a 5×10^6^ steps slab simulation of 300 LAF1 IDRs and 100 A15 RNAs. LAF1 IDRs are in green or blue, whereas RNAs are in red or purple color. (D) The last structure of a 2×10^6^ steps droplet simulation of 750 LAF1 IDRs and 250 A15 RNAs. The color scheme is the same as (C). (E) Radial distribution function of protein and RNA CG particles as a function of the distance from the center of mass of the LAF1 droplet (*r*). The structure of the condensation is shown as transparent background for reference.

In conjunction with the HPS model for protein IDR, the HPS model for RNA has also been recently developed [32]. The HPS RNA model has a different resolution from the other CG models described above, with one CG particle per nucleotide. Nevertheless, we also implemented this model in GENESIS, preparing for possible applications in the study of biophysical condensations involving both protein and RNA. Here we built and simulated two systems of IDR from the protein LAF1 and 15-nt (nucleotide) poly-A RNA (hereafter called A15). The two systems consisted of 300 LAF1 together with 100 A15 (57,300 particles in total) and 750 LAF1 together with 250 A15 (143,250 particles), respectively. The slab method was used to simulate the first system for 5 × 10^6^ steps, with the simulation box of 20*nm* × 20*nm* × 200*nm* (Fig 6C). The second system was simulated in a cubic box of size (100*nm*)^3^ for 2 × 10^6^ steps (Fig 6D). As can be seen in Fig 6C and 6D, in both situations, the A15 RNA entered the condensation of LAF1. We also quantitatively analyzed the co-condensation of LAF1 and RNA by plotting the radial distribution function (RDF) of protein and RNA particles as a function of the distance from the center of the LAF1 droplet (Fig 6E). There is a higher density of RNA inside the LAF1 condensation than in bulk (Fig 6E). These results are consistent with the previous simulation results [32].

### Application 3: virus capsid

We also explored the ability of our GENESIS CG implementation to simulate large-scale biological systems, such as a virus capsid. Here we reported our modeling and simulation of the herpes simplex virus (HSV) capsid, whose high-resolution structure has been recently solved with the cryo-electron microscopy (cryo-EM) (PDB entry: 5ZAP) [76]. The capsid comprises about 4000 proteins assembled into 12 pentons (pentameric blocks) and 150 hexons (hexameric blocks), and 320 triplexes gluing all the subunits [76]. The number of atoms and the length scale exceed the PDB format’s upper limit for such a vast structure. Therefore, the structural information of the HSV capsid was recorded in a file with the mmCIF format. We used the GENESIS-CG-tool to parse this file and generate the CG structure, which consists of 1,687,980 CG particles in total (Fig 7A). The AICG2+ potentials were used to model the whole structure. After generating the necessary input files, we then simulated the structure for 2 × 10^6^ steps. We tracked the radius-of-gyration (*R_g_*), the root-mean-square-deviation (RMSD), and the native-ness (*Q,* see Supplementary Methods for definition) of the whole structure (shown in Fig 7B). As can be seen, during our simulation, the structure was stably maintained. Although here we only show a short equilibrium simulation of the HSV capsid, to our knowledge, it is one of the largest biological systems simulated with the residue-level CG models. We expect to run CG simulations of similar systems using GENESIS to probe the conformational changes and dynamic assembly or disassembly of the capsid.

**Fig 7.**
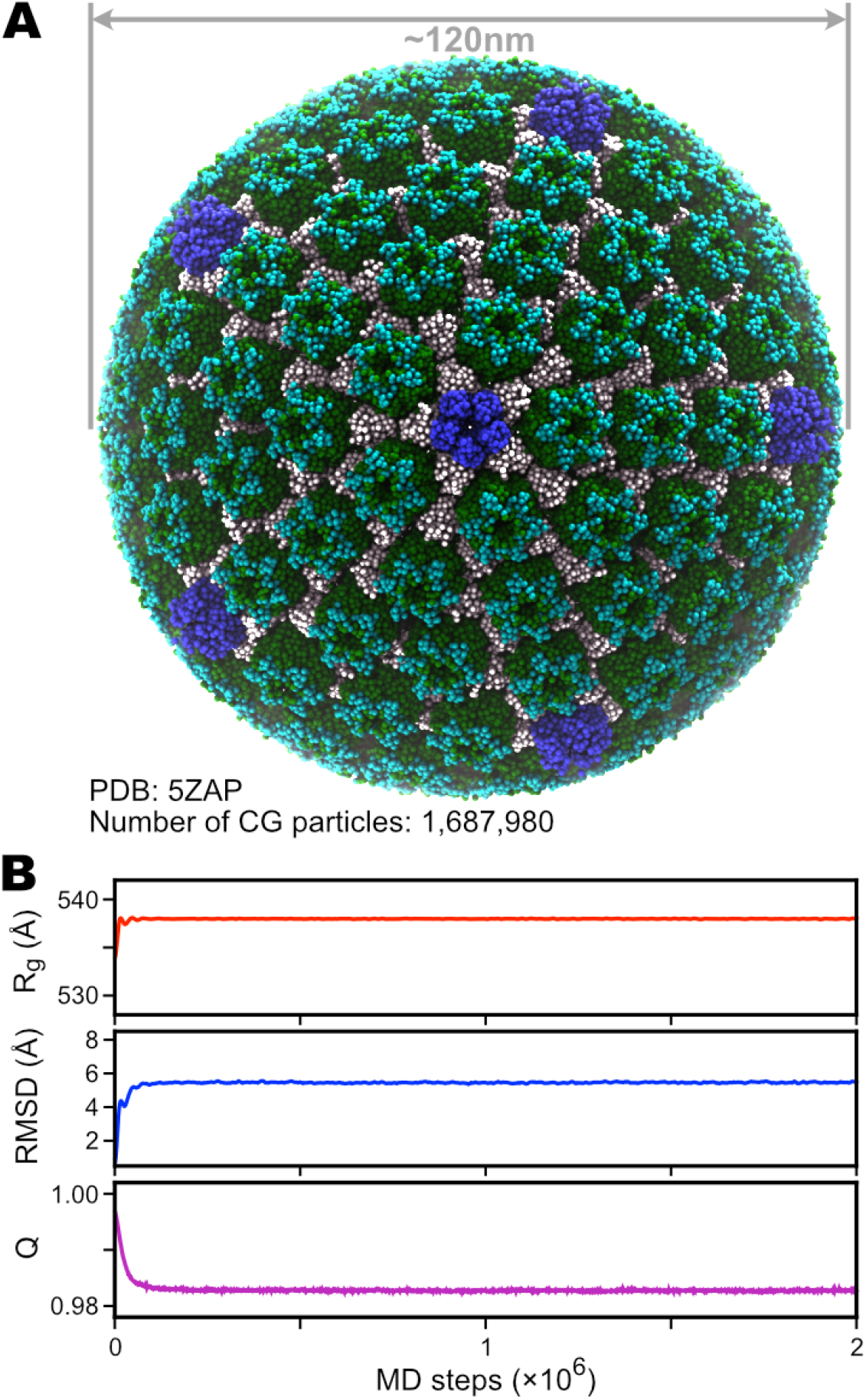
Simulation of the herpes simplex virus (HSV) type 2 B-capsid. (A) Structure of the coarse-grained HSV capsid, based on PDB entry 5ZAP. The pentameric blocks (pentons) are colored in blue, the hexameric blocks (hexons) are colored in green (VP5) and cyan (VP26). All the other proteins that glue together the pentons and hexons are colored in white. (B) Time series of the radius of gyration (*R_g_*), RMSD, and nativeness (*Q*) of the whole structure during a 2×10^6^-step simulation.

### Application 4: chromatin

Eukaryotic chromatin is the architecture that stores genetic information by packaging DNA into supercoiled structures around the histone octamer proteins. In addition to the function of genome organization, chromatin also hosts many biochemical reactions around the information flow between DNA and RNA. Due to its extraordinary importance, chromatin and related biological phenomena have been widely studied by MD simulations at different length scales and models at different resolutions [4,77–79]. In particular, residue-level CG models, including those we implemented here (*V_AICG2+_, V_3SPN.2C_, E_PWMcos_*, and *E_HB_*), have shown their abilities to decipher the dynamics of the nucleosome, which is the building block and basic unit of chromatin [25,26,56,57,80]. Here we tried to expand these residue-level CG MD simulations to a larger length scale. Instead of using a natural genomic sequence and constructing a Hi-C experiment-based structure, we chose poly-CG as the DNA sequence and built an artificial chromatin structure by connecting single nucleosome structures (based on PDB 1KX5) [68] with linear linker DNAs. The constructed structure contains 1024 nucleosomes linked by a 219,213-bp dsDNA (see Fig 8A and 8B). More details of the structure modeling can be found in the Supplementary Methods.

**Fig 8.**
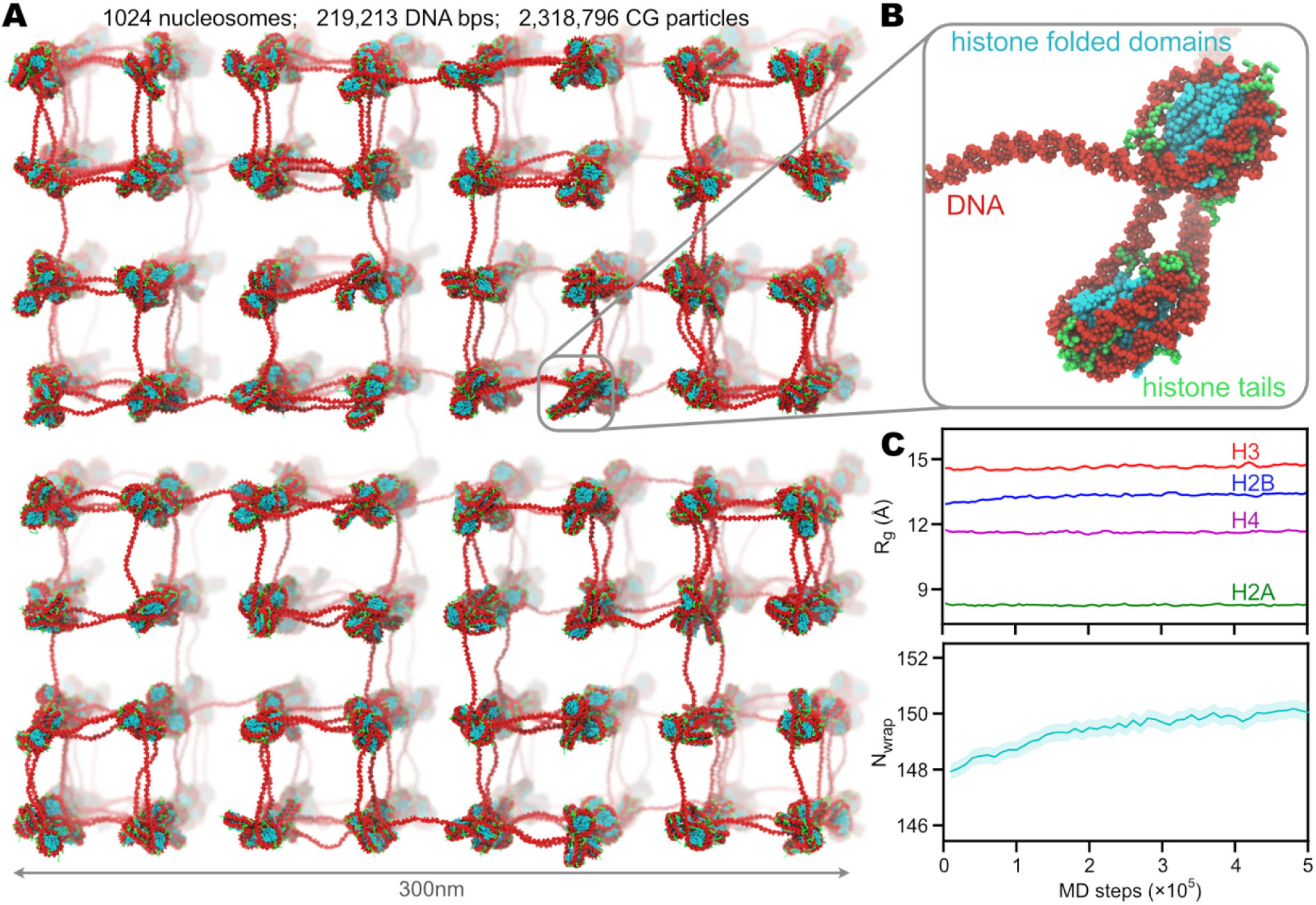
MD simulation of artificial chromatin composed of 1024 nucleosomes. (A) Structure of the whole artificial chromatin. DNA is colored in red, and histones are colored in cyan (for the folded domains) and green (tails). (B) A zoom-in structure of two adjacent nucleosomes as a part of the artificial chromatin. (C) Time series of the radius-of-gyration (*R_g_*) of the N-terminal tails of the histones (top) and the number of wrapped DNA base-pairs on histone (bottom). The solid lines represent the average values, and the shadowed regions show the standard errors.

Similar to previous studies [25,56], we used *V_AICG2+_* for histones, *V_3SPN.2C_* for DNA, and *E_HB_* for hydrogen bonds between histone and DNA phosphates. In addition, we applied *V_HPS_* to the flexible histone tails (see Supplementary Methods for definition). We then performed 5 × 10^5^-step MD simulations of this system at a temperature of 300K and 150mM ionic concentration. The time series of *R_g_* of the four types of N-terminal histone tails and the number of wrapped DNA base-pairs are shown in Fig 8C. These results were based on the statistics over the 1024 nucleosomes. This example illustrates the possibility of using our GENESIS CG implementation to carry out studies on the large-scale chromatin dynamics with accurate modeling of DNA and both well-folded and intrinsically disordered parts of proteins.

## Discussion

In this paper, we have implemented residue-level CG models of biomolecules in GENESIS MD software and have applied them in CG MD simulations of various biomolecules by combining different CG models. The combination of the AICG2+ model for proteins with the 3SPN.2C model for DNA and the PWMcos potential for protein-DNA sequence-specific interactions has been applied in the previous studies [57]. Whereas the combination of the HPS model for IDR and AICG2+ for folded protein is tested, for the first time, in this new framework. We also noticed several recently published CG models at an equivalent or similar resolution as ours, such as the TIS RNA model [18] and the iSoLF lipid model [81]. We will consider incorporating these models into GENESIS to cover a broader range of target biomolecules. In the applications presented here, we used the default interaction parameters in GENESIS. For more practical applications, such as genome-scale chromatin folding or phase-separated membrane-less condensations, one may need to more carefully optimize interaction terms between the IDR and the other components and their parameters. If necessary, the parameters should be calibrated carefully with experimental information as references. The GENESIS CG framework provides a convenient environment for these attempts.

Similar to the current work, other groups have also released combinatorial implementations of CG methods, such as the OpenAWSEM-Open3SPN2 within the OpenMM framework [38], the 3SPN.2C packages in the LAMMPS framework [33,34], and the CafeMol package [39]. Particularly, the OpenMM package utilized GPU calculations to gain better performances [38]. Compared with its implementation, our development aims to cover more models and provide a more flexible and feasible environment for the studies of larger-scale biomolecular systems. CafeMol has been successfully applied for CG simulations in various biomolecular studies [21,24,25,57,82,83]. However, it is not designed for the high-performance computing of larger-scale systems. We have optimized our code in GENESIS to enable high-performance simulations of systems consisting of millions of particles on regular computers (Figs 4, 7, and 8). There is an upper limit of particle number with the atomic decomposition scheme in atdyn, which essentially limits our the applicability with the current code to huge biological systems. The domain-decomposition strategy and the efficient load balancer suitable to residue-level CG models are required to solve this problem. These two will be implemented in a new MD engine in GENESIS.

Although the CG models have boosted the computational efficiency to be orders of magnitudes faster than the all-atom force fields [17], the speed-up still looks insufficient when studying systems of the organellar or cellular scales. It is necessary to employ enhanced sampling methods, many of which [84–87] have also been implemented in GENESIS. Taking this advantage, we will combine CG MD simulations with some of the enhanced sampling methods to obtain more statistically reliable data with reasonable computational times and to enable free-energy analysis in the organellar or cellular scales.

In parallel to the advances in computational biophysics, mesoscopic pictures of biomolecules in the organellar or cellular environments have been better described using the latest experimental techniques, such as cryo-electron microscopy (cryo-EM) [88], cryo-electron tomography (cryo-ET) [89], high-speed atomic force microscopy (AFM) [90], and in-cell NMR [91]. Integrating structural information by these experiments with computational biomolecular dynamics is one of the most important tasks in this research field. Recently, there have been several attempts [92–94] to make more realistic structural models of sub-cellular systems by utilizing available experimental results as much as possible. We hope that the implementation of residue-level CG models can provide more realistic structural dynamics of heterogeneous biological systems together with recent experimental information and modeling strategies.

For this purpose, multi-scale and multi-resolution simulations are helpful in the trade-off between modeling accuracy and computational efficiency, if we can switch the resolutions of target biological systems easily from this residue-level CG models to other CG models with different approximations (MARTINI [13], SPICA [95], PACE [96], etc.) or all-atom force field models (AMBER [97], CHARMM [98], GROMOS [99], etc.). In the GENESIS software, most of the above models are already available, and some of them are being implemented by us. In the near future, we may be able to investigate the inside of the cell at various resolutions, as Google Earth can do visualizations of both countries and local towns.

In summary, we have implemented several residue-level CG models in the GENESIS MD software, including the AICG2+ model for protein, the 3SPN.2C model for DNA, the PWMcos model for protein-DNA interactions, and the HPS/KH model for IDR and RNA. The MD input data for these CG models are given in a GROMACS-style file format by an easy-to-use toolbox, GENESIS-CG-tool. The computational performance of CG MD simulations is optimized to regular computers by preparing the nonbonded interaction lists suitable for the CG potential functions, vectorizing do-loop structures in the time-consuming subroutines, parallelizing the generation of random numbers in Langevin dynamics, and so on. With these attempts, GENESIS CG MD simulation is applied to large biological systems containing millions of CG particles efficiently on regular computers. The models and programs implemented here would be valuable for investigating heterogeneous biological phenomena involving folded/disordered proteins, RNAs, and DNAs at high concentrations. Implementations for larger systems containing billion CG particles and the use of enhanced sampling in the systems will follow the current implementation in the GENESIS software.

## Availability and future directions

The source code and user manual of GENESIS v1.7.0 is available at https://www.r-ccs.riken.jp/labs/cbrt/website, and GENESIS-CG-tool is both included as a part of GENESIS v1.7.0 and deployed at https://github.com/noinil/genesis_cg_tool. Tutorials of performing CG simulations with GENESIS v1.7.0 are open on https://www.r-ccs.riken.jp/labs/cbrt/tutorials2019/website. All the MD simulation files and data to produce the results are available from https://github.com/RikenSugitaLab/cg-development-GENESIS-1.7.0. We plan to use GENESIS v1.7.0 to study large-scale biological phenomina such as LLPS of protein and nucleic acids. We also plan to implement more CG models of biomolecules into GENESIS.

## Acknowledgments

The authors thank Ai Shinobu and Giovanni B Brandani for testing the developments of CG models in GENESIS v1.7.0.

## Supporting information

**Supplementary Methods: Supplementary information of molecular dynamics details.**

**Fig S1: Examples of using GENESIS-CG-tool to prepare CG topology and coordinate files.**

**Fig S2: Charge distribution on the surface residues of protein sry.**

